# Dedicated setup for the photoconversion of fluorescent dyes for functional electron microscopy

**DOI:** 10.1101/622639

**Authors:** Katharine L. Dobson, Carmel L. Howe, Yuri Nishimura, Vincenzo Marra

## Abstract

Here, we describe a cost-effective setup for targeted photoconversion of fluorescent signals into electron dense ones. This approach has offered invaluable insights in the morphology and function of fine neuronal structures. The technique relies on the localized oxidation of diaminobenzidine (DAB) mediated by excited fluorophores. This paper includes a detailed description of how to build a simple photoconversion setup that can increase reliability and throughput of this well-established technique. The system described here, is particularly well-suited for thick neuronal tissue, where light penetration and oxygen diffusion may be limiting DAB oxidation. To demonstrate the system, we use Correlative Light and Electron Microscopy (CLEM) to visualize functionally-labelled individual synaptic vesicles released onto an identified layer 5 neuron in an acute cortical slice. The setup significantly simplifies the photoconversion workflow, increasing the depth of photoillumination, improving the targeting of the region of interest and reducing the time required to process each individual samples. We have tested this setup extensively for the photoconversion of FM 1-43FX and Lucifer Yellow both excited at 473 nm. In principle, the system can be adapted to any dye or nanoparticle able to oxidize DAB when excited by a specific light wavelength.

## 1 Introduction

The study of presynaptic organisation requires the analysis of both functional and structural aspects of axons and their synaptic compartments (for a review see Debanne et al., 2011). While, for structurally well-characterized synapses, vesicular release can be studied using mainly electrophysiological methods (Jonas et al., 1993; Neher and Sakaba, 2001; Pulido et al., 2015), the study of central synapses often requires a combination of imaging methods. A number of superresolution techniques can provide information on the morphology of labelled structures with nanometer accuracy (see Gramlich and Klyachko, 2019), however, Electron Microscopy (EM) is still the only imaging technique able to provide contextual information at the ultrastructural level. Classic EM cannot provide any functional information and, to overcome this limitation, a number of Correlative Light and Electron Microscopy (CLEM) techniques have been developed over the years to study live tissue (Maranto, 1982; Harata et al., 2001; Shu et al., 2011; Darcy et al., 2006; Peddie et al., 2017; Ratnayaka et al., 2011; de Beer et al., 2018). Many CLEM techniques rely on the conversion of the fluorescent signal into an electron dense one and when this process requires light activation it is generally referred to as photoconversion. For over 30 years, photoconversion of diaminobenzidine (DAB) has been used to observe, at the ultra-structural level, functionally identified neurons (Maranto, 1982; Smith and Bolam, 1990; Viney et al., 2013). Harata et al. (2001) have shown for the first time that FM1-43 photoillumination in fixed cell cultures can be used to precipitate DAB into an osmiophilic polymer selectively in synaptic vesicles that underwent release and endocytosis in the presence of the dye. In spite of its usefulness, DAB photoconversion in thick neuronal tissue has only been employed by a relatively small number of research groups. A number of different protocols have been published to describe the technique in a range of different preparations and using different modes of light delivery to drive the photoconversion (Opazo and Rizzoli, 2010; Marra et al., 2014; Sabeva and Bykhovskaia, 2017).

Here, we present a low-cost system to drive targeted photoconversion of DAB in thick neuronal tissue. The intent of this article is not to describe the entire photoconversion procedure (already presented by Marra et al., 2014), but to provide instructions to build a photoconversion setup to easily identify and illuminate a region of interest in a controlled manner.

## 2 Equipment

The photoconversion setup (Figure 1) consists of a high-power excitation light laser, an optical microscope for monitoring the sample during photoconversion, a simple stage to hold the sample and supply the oxygen required for DAB polymerization. We have included a list of the components (Table 1) required to build the system described here but we have not included general laboratory supplies.

**Figure 1:**
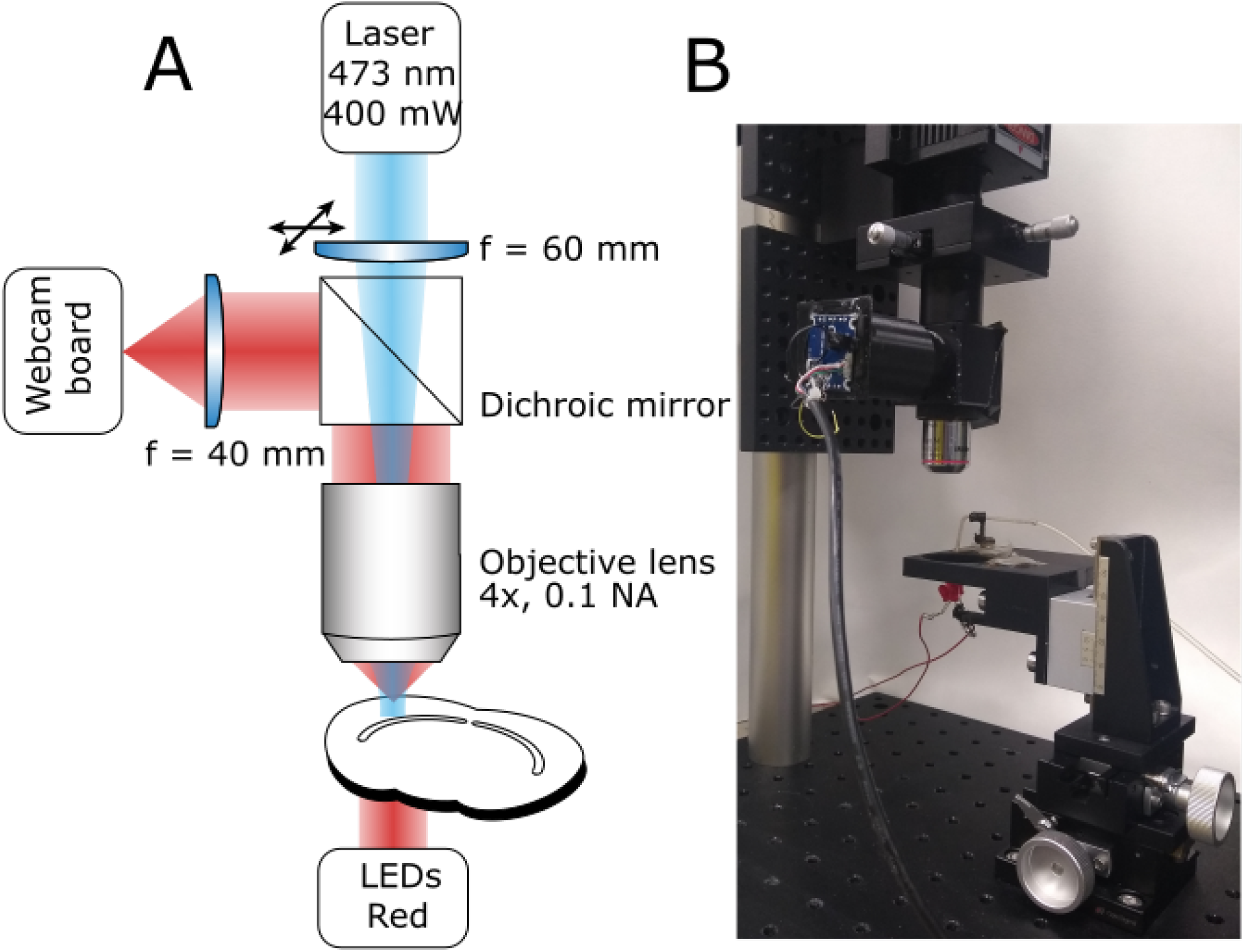
Photoconversion Rig. **A** Diagram of the light path for both sample illumination (blue) and visualization (red). **B** Photograph of a working setup including sample illumination and visualization paths and sample holder.

### 2.1 Excitation light path

The system’s main purpose is to deliver high-power excitation light over a large volume of tissue. The wavelength of the excitation light will depend on the fluorophore used. The system described here is built to photoconvert FM 1-43FX and Lucifer Yellow (excitation peaks at 473 nm and 425 nm, respectively), for this reason we chose a 473 nm Diode-Pumped Solid State (DPSS) laser as the light source, however, high-power (>300mW) single-wavelength LEDs would also work. The light path is extremely simple to minimize attenuation (Figure 1A), it consists of a lens focussing the laser beam on the back focal aperture of a 4X air objective. Underfilling the objective aperture produces an approximately collimated beam providing a larger volume of evenly photoilluminated tissue. The light will still be scattered by the tissue but the illuminated region will be effectively thicker driving the reaction over a larger volume for the same exposure time. Photoconverted vesicles can be observed at 85 μm below the slice surface, a 35 μm improvement over the previous method (Marra et al., 2012; data not shown). Between the focussing lens and the objective, the light goes through a 505 nm short-pass dichroic mirror. While the dichroic mirror attenuates the laser intensity, it also allows simultaneous visualisation of the sample and of the photoilluminated area for precise positioning. This system ensures that over 20% of the laser output reaches the focal plane of the objective with an homogeneously illuminated area of 0.65 *mm*^2^. The fine position of the illuminated area can be adjusted using an XY lens translator. This component is not strictly required, however, it allows fine adjustments improving accuracy and reducing the chance for experimenters’ error. Given the large volume illuminated, the power density is comparable to the previously reported one (Marra et al., 2014) and unlikely to directly damage fixed tissue. However, non-specific DAB oxidation may lead to precipitates that could obscure or damage the region of interest. The easiest solution to this issue, is to perform live labelling at least 20 μm below the surface of the tissue, when possible.

### 2.2 Sample visualization path

A critical step, in the photoconversion procedure, is the identification of the region of interest. In this photoconversion system, the objective used to deliver the excitation light is also used to image the sample which is illuminated from below with dim throughhole red LEDs. The red LEDs’ light is unlikely to excite and photobleach the fluorophores and can be used to monitor the progress of the photoconversion reaction. The red light illuminating the sample will reach the objective, be reflected by the dichroic mirror and imaged using a simple webcam (Figure 1). We found that a large field of view is useful to identify the region of interest, particularly in acute brain slices. Such large field of view can be obtained by using a camera with a large sensor (e.g. >1/1.8”) or by reducing the objective’s magnification. Here, we have used an inexpensive webcam with a small sensor 1/4”, removed the casing and glued it to a 3D-printed adaptor (Figure 2A). The lens creating the image on the webcam sensor is placed at only 70 mm from the objective’s back aperture, providing an effective magnification of 1.5X. This provides an adequate compromise between magnification and field of view. The ability to identify the region of interest with a large field of view greatly speeds up the positioning process down to a few minutes. Depending on user’s experience and region of interest’s size, the positioning process can take up to 45 minutes when using a high power objective and following a ‘roadmap’ acquired before tissue fixation, particularly without the aid of fluorescence.

**Figure 2:**
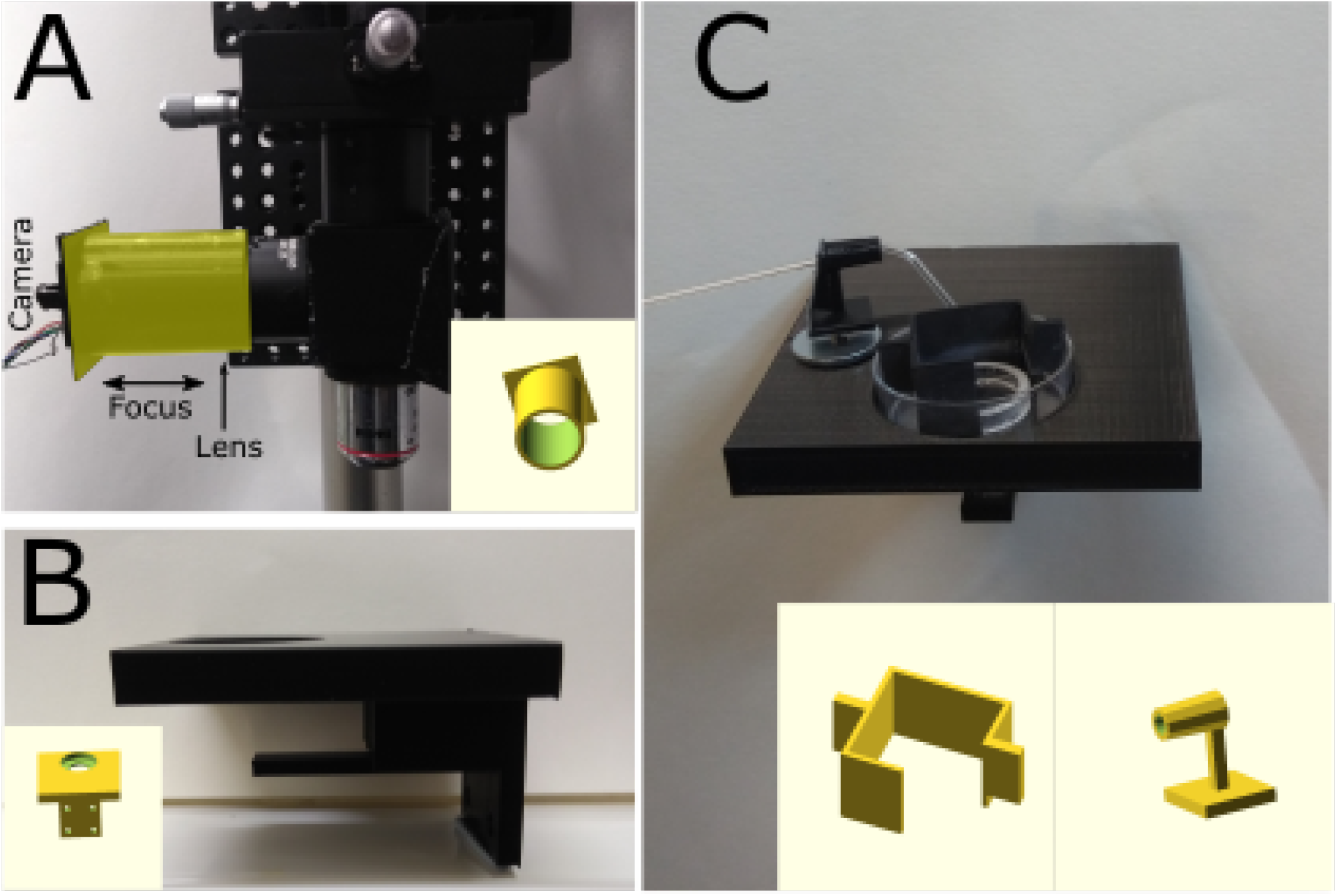
3D-printed components. **A** Photograph of a simple adaptor (highlighted in the yellow) for mounting a webcam board onto a Thorlabs 1” lens tube. The lens focussing the image on the sensor is placed inside the tube and the 3D-printed adaptor can be moved to focus the image. Insert: 3D model of the adaptor. **B** Photograph of stage mounted on 3-axis translator. Metal discs were glued and covered with transparent plastic to avoid DAB-induced oxidation. Insert: 3D model of the stage. **C** Sample in a 35mm petri dish held in position by a 19mm glass ring with nylon string across its middle and plastic box that prevents sample movement and bubbles covering the sample.The sample would placed under the glass ring. Glass capillary holder with a magnet mounted at its bottom to hold it in position. The capillary is used to oxygenate the solution and facilitate DAB photoconversion. Insert: 3D model of petri dish box and capillary holder.

### 2.3 Stage and oxygen delivery

To minimize the cost, a stage was designed using OpenSCAD (openscad.org) and 3D-printed using an Ultimaker2^+^ 3D-printer. The stage is connected to the 3-axis translator and used to hold and position the sample. Magnetic metal should be glued on the stage to position a capillary holder (Figure 2C). The stage includes a support for the throughhole LEDs used to illuminate the sample (Figures 1B and 2B). Once ready for photoconversion, the fixed tissue is placed in a 35 mm petri dish containing a plastic insert (Figure 2C) and held down with a glass O-ring (19 mm O.D.) with nylon strings (Marra et al., 2014). Once the user has identified the region of interest, PBS can be replaced with the DAB solution and oxygenated using Carbogen (95% O_2_ and 5% CO_2_). Carbogen is delivered using a glass capillary with an opening of >150 μm. The capillary is held by a 3D-printed holder (Figure 2C) with a magnet glued to its base for positioning. The plastic insert has two functions: keeping the sample and the O-ring in the center of the petri dish and avoiding bubbles from floating over the sample and thus scattering the light (Figure 2C).

## 3 Materials and Methods

### 3.1 Brain slice preparation

This study was conducted in accordance with the UK Animals (Scientific Procedures) Act, 1986 and following institutional regulations by the University of Leicester, animals were sacrificed under Schedule 1 and the project did not require full review and approval by the Home Office. Experiments were performed in CBA/Ca mice of either sex postnatal day 15—20. Animals were humanely killed by cervical dislocation and cessation of circulation was confirmed by severing the carotid artery. The brain was rapidly removed into ice-cold oxygenated artificial cerebrospinal fluid (aCSF) consisting of (in mM): 125 NaCl, 2.5 KCl, 26 NaHCO_3_, 1.25 NaH_2_PO_4_, 25 glucose, 1 MgCl_2_, 2 CaCl_2_; pH 7.3 when bubbled with 95% O_2_ and 5% CO_2_. Auditory cortex slices were cut at a thickness of 350 μm using a vibrating microtome (Leica VT1200S) at an angle of 15° off horizontal to preserve thalamocortical projections (Llano et al., 2014). Slices were allowed to recover for 20 minutes at 32°C before being cooled to room temperature.

### 3.2 Electrophysiology and cell labelling

Whole-cell recordings were made from visually identified layer 5 pyramidal neurons. Intracellular recording solution consisted of (in mM): 115 KMeSO_4_, 5 KCl, 10 HEPES, 10 creatine phosphate, 0.5 EGTA, 2 MgATP, 2 Na_2_ATP, 0.3 Na_2_GTP, 290 mOsm, with the addition of Lucifer Yellow (1 mg/mL, Sigma-Aldrich, ex/em 425/540 nm). After 5 minutes in whole-cell configuration a Z-stack image of the dye-loaded cell was acquired for later reference. A glass stimulating electrode filled with FM 1-43FX (20 μM in aCSF) (Thermofisher) was positioned ~ 150 μm from the recorded cell soma at a depth equal to that of the recording electrode. Prior to the onset of dye application the stimulus intensity was set to give a reliable excitatory post-synaptic current (EPSC) amplitude of ~200 pA, to ensure the recruitment of a large number of presynaptic terminals.

### 3.3 Vesicle labelling

FM 1-43FX (Thermofisher, ex/em 473/585 nm) dye was locally applied by providing 1.03 bars of positive pressure for a total of 5 minutes. Two minutes after the start of dye application presynaptic terminals were stimulated at 10 Hz for 60 seconds using a stimulus isolator (Digitimer). Dye application continued for further 2 minutes for a total of 5 minutes dye application and 10 minutes in whole-cell configuration. The recording electrode was then carefully withdrawn to reseal the recorded cell and minimize Lucifer Yellow leakage.

### 3.4 Fixation and photoconversion

Fixation and photoconversion were performed ensuring the samples’ exposure to light was kept to a minimum and following the protocol of Marra et al. (2014). Slices were carefully removed from the recording chamber and fixed in 6% glutaraldehyde/2% paraformaldehyde in PBS using microwave-assisted fixation. Samples were blocked in 100 mM glycine solution for 1 hour, rinsed in 100 mM ammonium chloride for 1 minute, then washed in PBS. The sample was then positioned on the photoconversion setup (Figure 1B) with a fine bore glass capillary immersed to allow continuous bubbling with Carbogen throughout. PBS was replaced with 1 mg/mL DAB solution prepared in oxygenated PBS and incubated in the dark for 10 minutes. This was replaced with fresh oxygenated DAB solution and the photoconversion reaction initiated using the 473nm DPSS laser (Figure 1A). Before starting the photoconversion reaction, the laser should be set to a low power setting and positioned on the region of interest using the fine XY lens translator while the transmission LEDs are on, to obtain an unequivocal image of the photoilluminated region on the sample. Reactions were monitored every 5–10 minutes and terminated when a defined dark region was visible on the slice. Since a number of factors may affect the time required to obtain a successful photoconversion, we recommend monitoring the process at regular intervals (~5 minutes) by powering down the laser and measuring the intensity of the transmitted light. In the first 10 minutes there should be a clear decrease in transmitted light intensity that will plateau when the photoconversion process is complete (see Marra et al., 2014). The photoconversion of FM-stained dead tissue will reduce light penetration as the reaction progresses. Using an approximately collimated beam ensures that a large volume is photoconverted before light penetration becomes an issue. The sample was then washed 3 × 5 minutes in PBS. An image of the photoconverted slice was captured and the sample trimmed for further processing.

### 3.5 Sample processing for electron microscopy

Processing of the fixed photoconverted tissue for electron microscopy follows the protocol of Deerinck et al. (2010). Briefly, samples were washed with 0.15 M sodium cacodylate buffer prior to osmication (2% OsO_4_, 1.5% K4Fe(CN)_6_). Samples were then washed with dH_2_O and incubated in 1% thiocarbohydrazide solution. Tissues were rinsed again in dH_2_O and thereafter placed in 2% osmium tetroxide. Following this second exposure to osmium the tissues were washed with dH_2_O then placed in 1% uranyl acetate at 4° overnight. As an optional step, samples were then washed again with dH_2_O and stained *en bloc* with freshly prepared 20 mM lead aspartate. After further dH_2_O washes samples are serially dehydrated using a gradient of ethanol solutions and finally with 100% acetone. Samples were infiltrated with Durcupan™ resin before being transferred to a BEEM capsule for embedding and polymerisation. With reference to the image of the photoconverted tissue the sample was trimmed, ultrathin (~70 nm) sections cut and mounted on copper grids. Sections were imaged on a JEOL 1400 transmission electron microscope equipped with an OptiMos (QImaging).

## 4 Anticipated Results

### 4.1 Simultaneous photoconversion of FM 1-43FX and Lucifer Yellow

To show potential applications of the photoconversion setup, we have combined electrophysiological recordings of a neuron filled with Lucifer Yellow via a patch pipette with simultaneous loading of synaptic vesicles with FM 1-43FX. FM-loading is obtained by puffing the dye with a glass pipette which is also used to deliver the stimulation required for the labelling of synaptic vesicles (10 Hz, 60 seconds), vesicular release is monitored recording the post-synaptic neuron in voltage clamp (Figure 3A). Importantly, the choice of stimulation frequency and duration will depend on the biological question asked. For example, a 1 Hz stimulation for 30 seconds can be used to study single site release probability (Branco et al., 2010), stimulation frequencies of 10 Hz or above for 1 or 2 minutes may be used to label the entire recycling pool (Fornasiero et al., 2012) and higher frequency stimulations for few seconds may be used to label the readily releasable pool (Rey et al., 2015). The stimulation frequency used will also depend on type of synapse probed. For example, at large release sites such as the Calyx of Held, a giant synapse in the auditory pathway, stimulations > 100 Hz are appropriate to recruit the total recycling pool (Lucas et al., 2018); similarly at neuromuscular junctions long high frequency stimulations (30 Hz, 5 minutes) may be needed (Denker et al., 2009).

**Figure 3:**
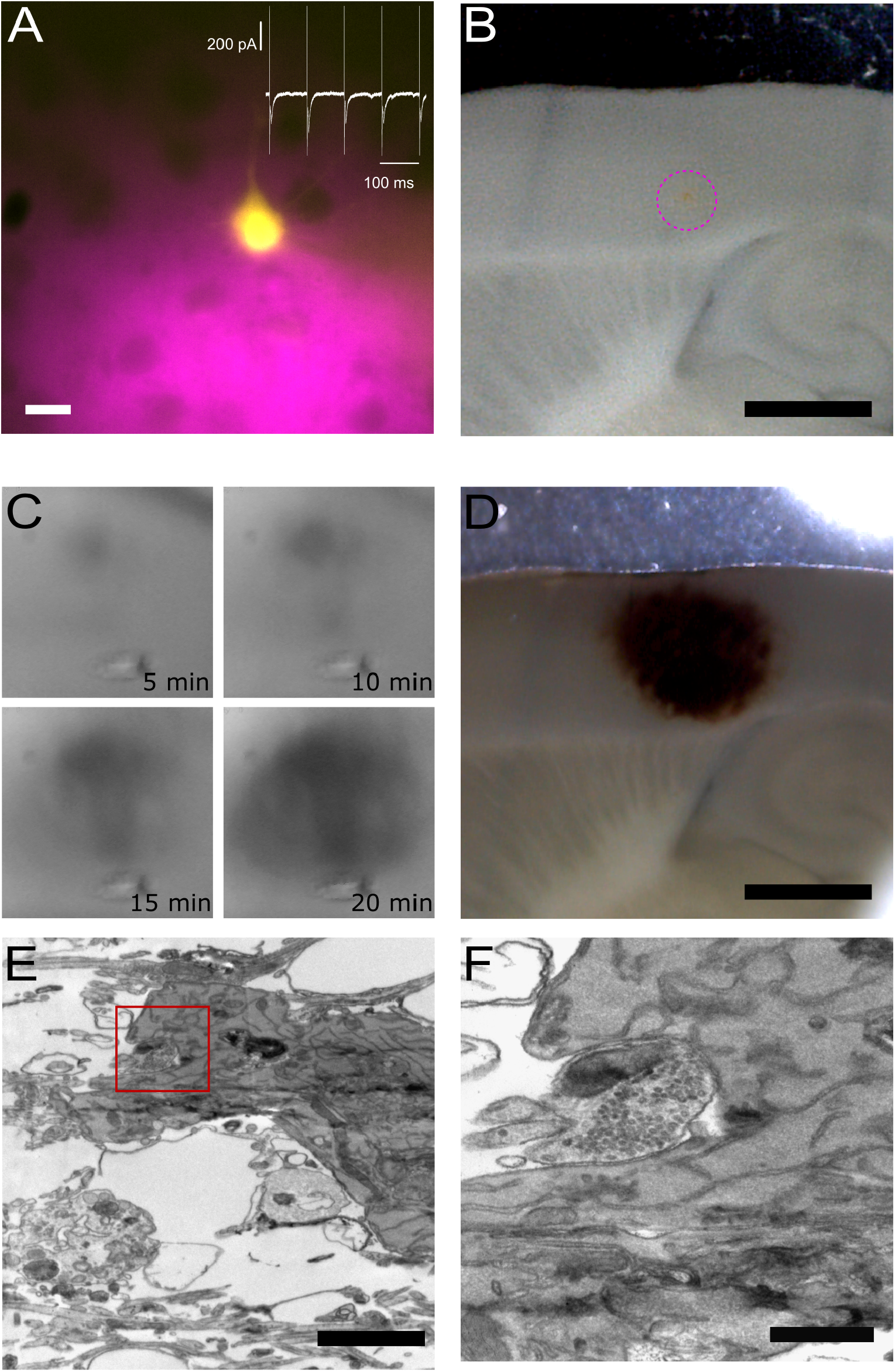
Pre- and post-synaptic photoconversion anticipated results. **A** Pyramidal neuron loaded with Lucifer Yellow with region of pressure-applied FM dye highlighted in magenta. Insert: whole-cell voltage clamp recording from loaded cell. Scale bar: 20 μm. **B** Acute slice following fixation, with FM dye region clearly visible (magenta dashed circle). Scale bar: 1 mm. **C** Development of the photoconversion product monitored at 5 minute intervals. **D** Photoconverted region extending across FM dye region and Lucifer Yellow-loaded neuron. Scale bar: 1 mm. **E** Electron micrograph of Lucifer Yellow-loaded neuron. Scale bar: 2 μm. **F** Higher magnification electron micrograph of the same spine as E, with FM-loaded vesicles observed in the presynaptic terminal. Scale bar: 500 nm.

Figure 3 provides an overview of typical results plus examples of the tissue at each stage of processing. The location of the two fluorophores, FM 1-43FX to label presynaptic vesicles and Lucifer Yellow to fill the post-synaptic cell, is visible throughout processing, thus giving the user confidence in the final structural information correlating with the functional readout from the electrophysiological assay. The site of delivery for FM 1-43FX can be easily visualized at low magnification (Figure 3B). Following fixation, the sample is moved to the photoconversion setup where the reaction’s progression can be observed as a darkening of the tissue (Marra et al., 2014). Before processing for electron microscopy the photoconverted region appears as a dark spot on the tissue (Figure 3D). Electron dense structures are readily identifiable in the resultant electron micrograph as a soma containing DAB precipitate (Figure 3E) and photoconverted vesicles observed in the terminal opposite the labelled postsynaptic neuron (Figure 3F). Lucifer Yellow’s absorption is only 25% at 473 nm and this appears sufficient to drive DAB photoconversion without excessive DAB-damage (Figure 3E).

Depending on the stimulation protocol, the approach demonstrated here can be used to investigate a number of key synaptic and cellular parameters, e.g. synaptic sites, release probability, recycling pool organisation, and in general the impact of pre- and post-synaptic structures on synaptic function.

## 5 Discussion

The functional study of axonal fine structures often requires a combination of fluorescence and electron microscopy. These measurements can be performed in parallel (Clayton et al., 2008; Chung et al., 2010; Cheung et al., 2010; Ratnayaka et al., 2012) or in a correlative way following the same axons and synapses from light to electron microscopy (reviewed by Branco and Staras, 2009; Begemann and Galic, 2016). In both cases, however, the measurements are limited to dissociated cell cultures. A number of new tools are being developed by commercial and academic research laboratories to simplify the CLEM workflow and to extend it to thick neuronal tissue. Some laboratories are developing new fluorescent proteins able to oxidize DAB in a light-independent way (Shu et al., 2011; Liss et al., 2015; de Beer et al., 2018), while others focus on new microscopes to improve the reliability of CLEM workflow (see Brama et al., 2016).

Here, we presented a simple setup to photoconvert fluorescent signals into electron dense ones. The setup is built to facilitate tracing of the region of interest in thick neuronal tissue improving CLEM workflows for *ex vivo* tissue. A dedicated setup can improve photoconversion’s reliability and throughput, the two main limiting factors in the diffusion of the technique. Compared with the use of standard epifluorescence microscope, our setup increases the depth of photoillumination, improves the targeting of the region of interest and reduces the time required to process each individual sample. Commercial epifluorescence microscopes have optics to ensure Köhler illumination of the sample, while offering a great advantage for imaging, such optics can attenuate a single wavelength light of ~90% before it reaches a dichroic mirror and filter, with further light attenuation. Additionally, most photoillumination approaches for thick tissue require the ‘sacrifice’ of a high power dipping objective to deliver light (Marra et al., 2014) as, once used for DAB photoconversion, a dipping objective should not be used for live tissue imaging. Additionally, air objectives will not be stained by repeated exposure to DAB, while dipping objectives, even with careful cleaning, will stain over time increasing the photoconversion time and affecting the quality of the image used for positioning.

The budget used for building the setup in 2016 was ~4500€, with over 50% of the budget used to purchase the laser. However, 473nm DPSS laser can now be found for much lower i prices or substituted by high-power single-wavelength LEDs, lowering the cost of the entire t· system to ~3000Є, comparable with the cost of a dedicated high quality dipping objective for photoconversion of thick neuronal tissue (Marra et al., 2014). We demonstrate that the setup described can be used for the simultaneous photoconversion of FM 1-43FX and Lucifer Yellow, allowing the study of cellular ultrastructure and activity and release probability at individual release sites.

## Supporting information

3Dprinted parts

## 6 Acknowledgements

The work was funded by the Wellcome Trust Seed Awards in Science (108201/Z/15/Z). The authors would like to thank the Core Bioscience Services for access to the experimental animals, the fluorescence microscope setup by Professor Nicholas Hartell and the electron microscope. In particular, we would like to thank Natalie Allcock for the assistance with the preparation of the samples for electron microscopy and the acquisition of the images. We would also like to thank Amy Richardson for building the PWM laser power regulator and Drs James McCutcheon and Joern Steinert for lending us components to test the early configuration of the setup.

## 7 Author Contributions Statement

KD acquired the data and tested the setup. YN tested the setup. CLH assisted with the design of the setup. VM designed, built and tested the setup. All authors contributed to the manuscript.

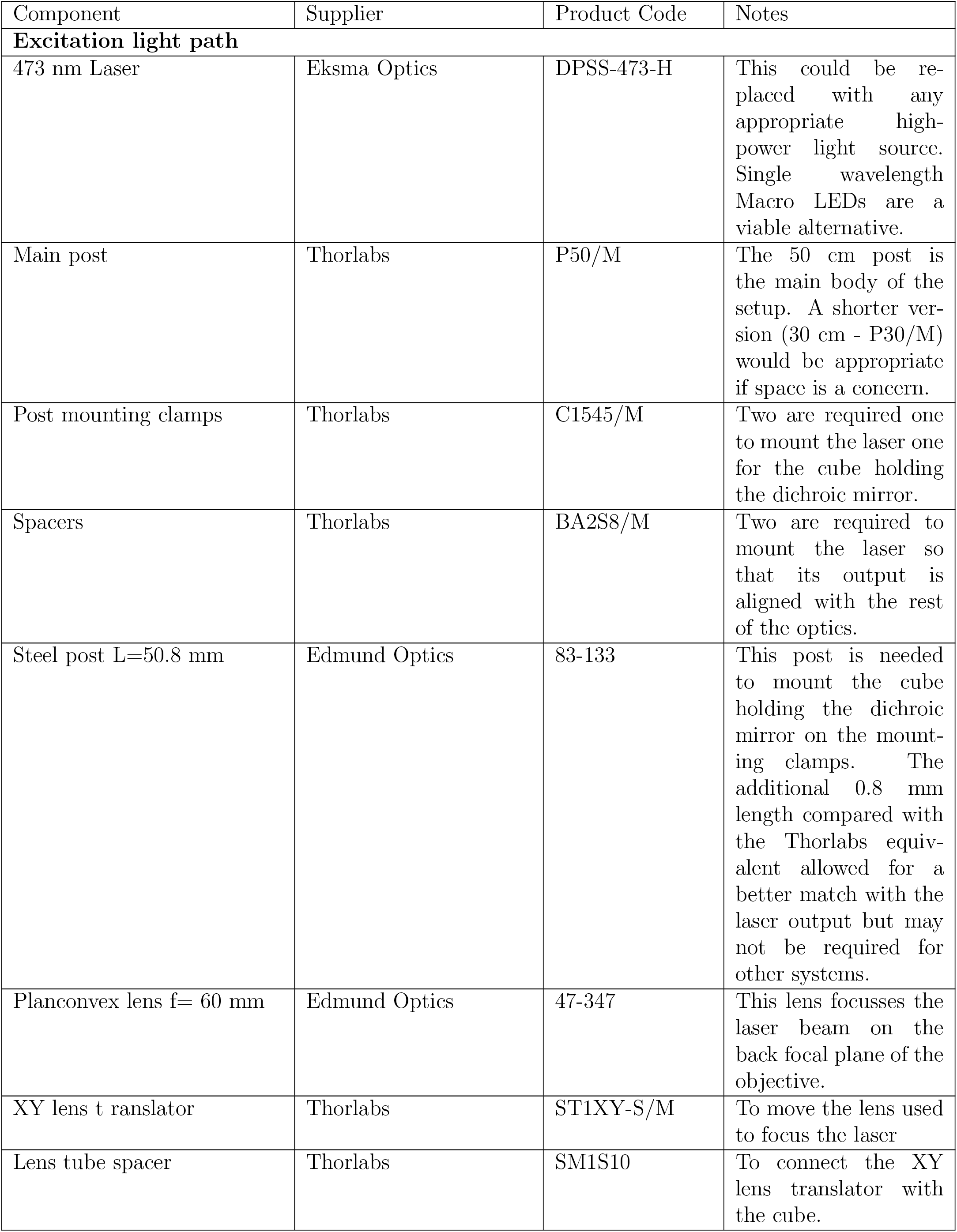
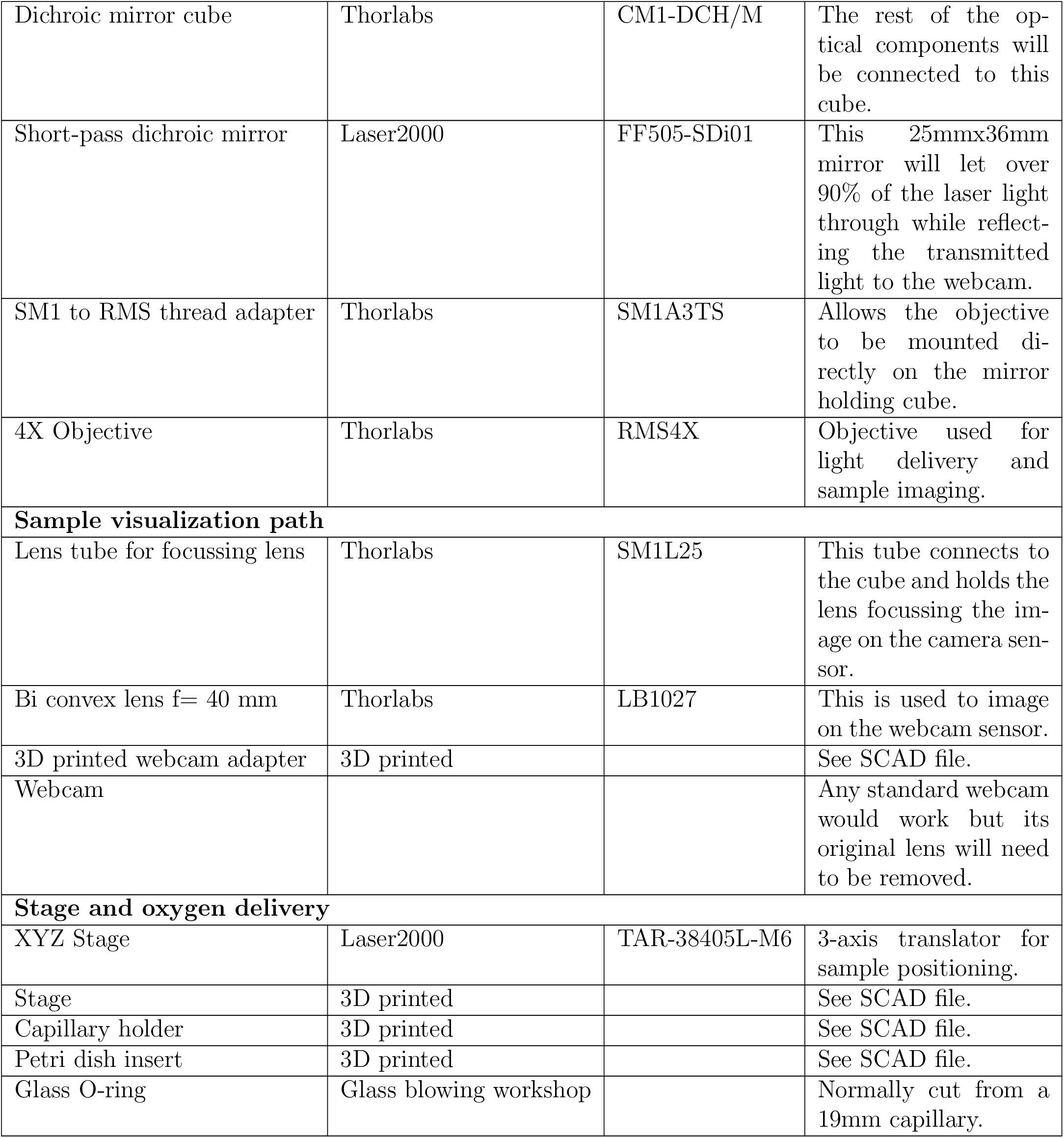

## Notes

#### Summary of Updates

Corrected text

https://github.com/enzomarra/PCrig

